# Deep Learning-Based 3D Leukocyte Differentiation Using Label-Free Higher Harmonic Generation Microscopy

**DOI:** 10.1101/2025.04.24.650417

**Authors:** Mengyao Zhou, Patrick José González, Tamara Dekker, Shiqi Zhang, Leonoor S. Boers, Hélène B. van den Heuvel, Annemiek Dijkhuis, Iris A. Simons, Jan Willem Duitman, Marie Louise Groot

**Affiliations:** Faculty of Science, Department of Physics, Laserlab, Vrije Universiteit Amsterdam, De Boelelaan 1105, 1081HV, Amsterdam, The Netherlands; Department of Experimental Immunology, Amsterdam UMC, location University of Amsterdam, Amsterdam, The Netherlands; Amsterdam Infection & Immunity, Inflammatory Diseases, Amsterdam, The Netherlands; Intensive Care Medicine, Amsterdam UMC, location University of Amsterdam, Amsterdam, The Netherlands; Department of Pulmonary Medicine, Amsterdam UMC, location University of Amsterdam, Meibergdreef 9, Amsterdam, The Netherlands

**Keywords:** Broncho-alveolar lavage fluid (BALF), blood fraction, deep learning, higher harmonic generation microscopy (HHGM), leukocyte differentiation

## Abstract

Both in clinical practice and translational research, cell differentiation of leukocytes provides important information used for diagnostics or insights into pathophysiological mechanisms. The current gold-standard method for bronchoalveolar lavage fluid (BALF) analysis involves histochemical staining of cytospins, followed by manual morphological quantification. However, this approach is labor-intensive, time-consuming, and highly operator-dependent, limiting its efficiency and throughput. This study proposes a deep learning framework for rapid, automated 3D leukocyte differentiation using label-free higher harmonic generation microscopy (HHGM). 3D leukocyte characterization was performed with label-free HHGM. Two deep learning models, ResNet 3D-50 and Vision Transformer (ViT) 3D, were trained, validated and tested for leucocyte differentiation on both BALF and blood fraction samples from 14 interstitial lung disease (ILDs) and 12 acute respiratory distress syndrome (ARDS) patients. Deep-learning model-prediction and cytospin analysis were performed by separate investigators. Results were compared using Bland-Altman analysis. The deep learning algorithm achieved >96% accuracy in quantifying neutrophils, eosinophils, lymphocytes, and macrophages/monocytes. Bland-Altman analysis showed mean differences of less than 3% between cytospin analysis and the deep learning based approach across all cell types. By integrating the label-free imaging capabilities of HHGM with deep learning, this study established a fast, accurate and high-throughput leukocyte differentiation in fresh BALF and blood samples. By significantly improving efficiency and reproducibility, this technology has the potential to transform clinical workflows and advance precision medicine.

## Background

Determining the leukocyte composition of bronchoalveolar lavage fluid (BALF) and blood is critical for diagnosing and managing various lung diseases [1-3]. For example, in acute respiratory distress syndrome (ARDS), neutrophils are the majority cells observed in both blood and BALF, reflecting the intense inflammatory response characteristic of the disease [1]. In interstitial lung diseases (ILDs), differential cell counts in BALF are used to guide the differential diagnoses of ILD. An abnormally high number of neutrophils and/or eosinophils, often referred to as “alveolitis”, are believed to reflect a lower respiratory tract inflammation [2, 4], whereas a high lymphocyte count suggests an ILD associated with granuloma formation, such as hypersensitivity pneumonitis or sarcoidosis [4]. Moreover, in eosinophilic asthma, increased eosinophil counts are frequently observed in both blood and BALF, contributing to airway inflammation and hypersensitivity, which are key features of the condition [3, 5].

Cell differentiation using histochemical staining of cytospin followed by manual microscopic examination is currently regarded as the gold standard for leukocyte analysis due to its ability to provide detailed morphological analyses [6, 7]. However, this approach is highly labor-intensive, time-consuming, and requires well-trained personnel. As a result, outcomes may vary significantly based on the observer’s expertise and experience. Flow cytometry is an alternative way to automatically differentiate leukocytes with high accuracy, but it generally cannot provide detailed cellular information and relies on labeling with multiple antibodies against specific cell markers to achieve high classification accuracy [8-10]. Despite being accepted by experts[8, 11, 12], this technique involves substantial costs for equipment (flow cytometer) and sample preparation materials (i.e., multiple antibodies). Higher harmonic generation microscope (HHGM), which integrates third harmonic generation (THG) and multiphoton excited autofluorescence (MPEF) microscopy, shows great potential for bridging this gap as it features 3D label-free and real-time imaging capabilities [13-15]. Applied to BALF and blood samples, this approach offers distinct advantages, such as minimal sample preparation, cost efficiency, and high throughput.

THG is sensitive to local variations in third-order nonlinear susceptibility, refractive index, and dispersion. Its signal is enhanced at interfaces and optical inhomogeneities that are similar in size to the beam focus [16, 17]. Due to the nonlinear dependence of signal generation on laser intensity, THG is confined to the focal region, giving it intrinsic axial optical sectioning capability. It enables near-complete visualization of tissue morphology, particularly excelling in imaging water-lipid interfaces, such as cell membranes and lipid-rich components [18]. MPEF, such as two-photon excited autofluorescence (2PEF) imaging, can detect mitochondria-related metabolic coenzymes such as NADH and FAD in various pathological conditions, including precancers [19]. Additionally, three-photon excited autofluorescence (3PEF) imaging can detect serotonin and NADH [20]. These modalities can be integrated and generate signals simultaneously, providing comprehensive subcellular information[21, 22].

For the resulting high-throughput and high-content images, equally fast and automated image analysis tools are required. There are deep learning frameworks that work well for identifying labelled cells in 2D sections [23-28]. However, 2D analyses on depth sectioned images may introduce bias as the number of detected cells can be either underestimated or overestimated depending on the depth sampling, and their size may be underestimated. Furthermore, 2D depth-sectioned images provide only partial information about the cell, potentially leading to errors in cell identification. Therefore, a 3D analysis of 3D volume measurements is required. When it comes to 3D cell analyses, two main challenges arise. First, annotating large numbers of cells is painstaking and labor-intensive. Second, the complex architecture of deep learning neural networks for big data, demand substantial computational resources to train and process images within a reasonable timeframe.

Therefore, in this study, we developed a computational pipeline in which ResNet 3D-50 and Vision transformer (ViT) 3D were trained to autonomously quantify the cell populations (neutrophils, eosinophils, lymphocytes, macrophages/monocytes) in fresh, label-free BALF and blood fraction samples without human intervention. This approach was validated and tested on BALF and blood fraction samples from ILD and ARDS patients, collected during routine clinical practice.

## Methods

### Sample collection and preparation

BALF and blood fraction samples were collected for clinical/diagnostic purposes from patients with ARDS and ILD, each following specific collection protocols, described below.

In ARDS patients admitted to the ICU at both locations of the Amsterdam UMC (AMC and VUMC) who exhibited no respiratory improvement, routine diagnostic bronchoscopy with BALF was performed according to standardized protocols. The decision to perform BAL was made by joint expert opinion during multidisciplinary team meetings. During bronchoscopy, the bronchoscope was wedged into a subsegment of a lung lobe at a site identified by high-resolution computed tomography (HRCT) or chest CT imaging. Four times 20 mL 0.9% NaCl were instilled into a single lung segment, with the recovered fractions collected separately. One milliliter of the first two fractions was reserved for label-free imaging. Additionally, venous blood was collected into heparinized tubes for blood fraction analysis. Blood samples were only available from ARDS patients.

For ILD patients, BALF was collected by instilling eight times 20 mL 0.9% NaCl followed by gentle aspiration. One milliliter of the final fraction was allocated for label-free imaging.

Following collection (Fig. 1A), BALF samples were processed following standard cytology protocols at the diagnostic lab of Amsterdam UMC (location AMC), resulting in Diff-Quick (DQ, RAL Diagnostics) stained cytospin slides (Fig. 1C). In every sample, at least 500 cells were evaluated and classified as macrophages, lymphocytes, neutrophils, eosinophils, or other by an experienced labtechnician using a standard light microscope at 40× magnification.

**Figure 1.**
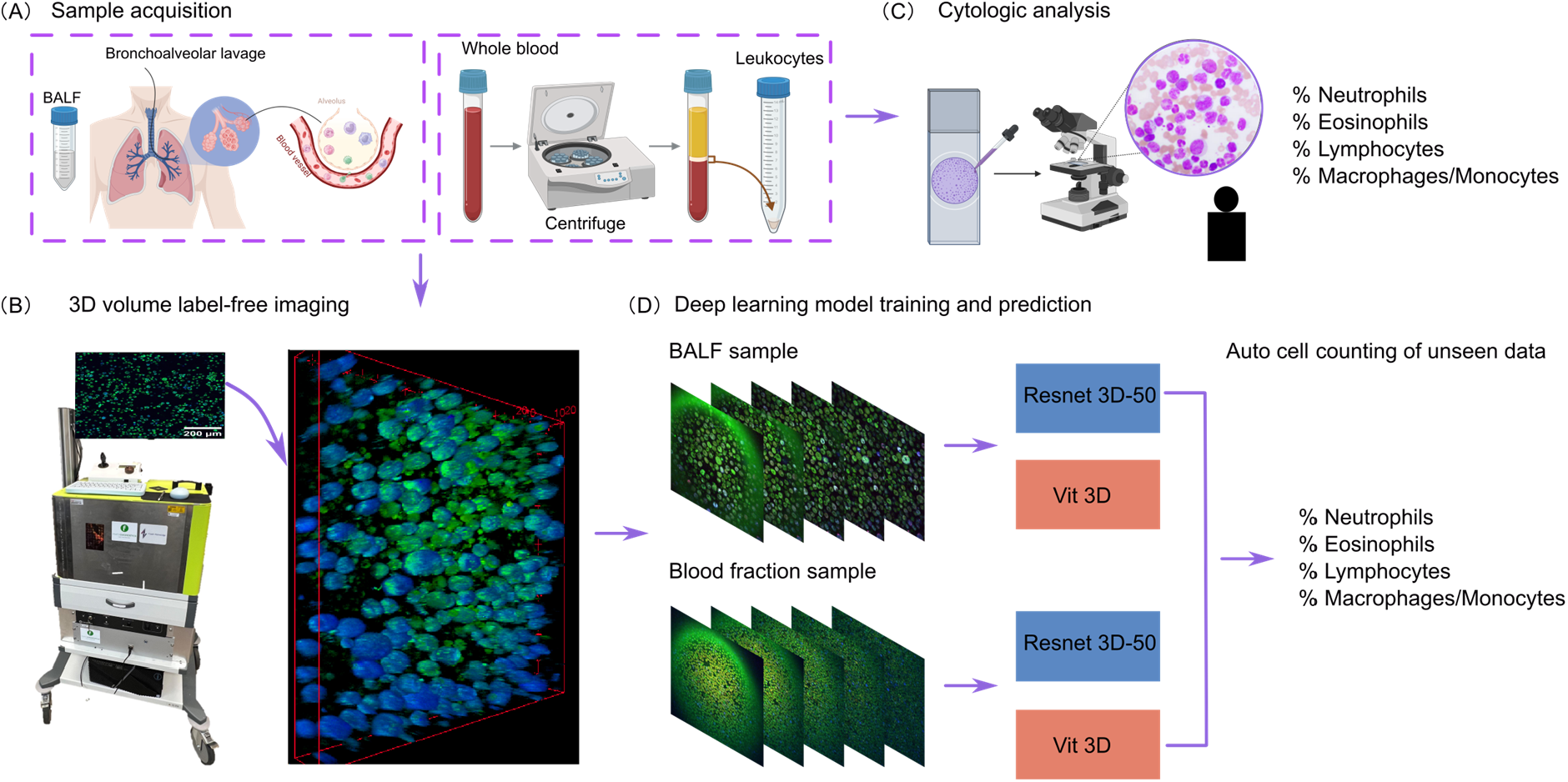
Diagrammatic illustration of overall workflow. (A) Sample acquisition of BALF and blood fraction sample. (B) 3D volume label-free imaging using HHGM. (C) Standard cytospin processing with DQ staining. (D) Training and prediction using deep learning models.

Blood samples were treated with erylysis buffer to lyse all erythrocytes, washed, and counted using a Coulter DxH500 hematology analyzer (Beckman Coulter, Brea, USA). Cell differentiation data for blood cells were obtained from the results, classified as monocytes, lymphocytes, neutrophils, eosinophils or other.

In parallel, 1 ml of each sample, containing approximately 0.5 million cells, were transported on ice to our lab (Imaging Center at Amsterdam UMC, location VUMC) for 3D label-free imaging, which was performed within half an hour after arrival. The sample was pipetted to a petri-dish (μ-Dish 35 mm, high, Ibidi GmbH) and placed in a portable HHGM for 3D volume label-free imaging (Fig. 1B).

In total, we collected 30 BALF samples and 16 blood fraction samples from 14 ILD patients and 12 ARDS patients. The characteristics of the patients and samples are shown in Table 1.

**Table 1.** Patient Characteristics.

|  |  | ILD (N=14) | ARDS (N=12) |
| --- | --- | --- | --- |
| Sample type |  | BALF | BALF and blood |
| Age, mean (minimum - maximum) |  | 65.8 (47-81) | 43.1 (19 -77) |
| Sex | Female, n (%) | 7 (50) | 3 (25) |
|  | Male, n (%) | 7 (50) | 9 (75) |
| Smoking status | Never, n (%) | 5 (36) | 4 (34) |
|  | Former, n (%) | 1 (7) | 1 (8) |
|  | Current, n (%) | 6 (43) | 6 (50) |
|  | Unknow, n (%) | 2 (14) | 1 (8) |

### HHGM setup

Figure 1 B shows the HHGM setup (co-developed with Flash Pathology B.V.) and a 3D rendering of a BALF sample. In this study, the microscope was upgraded to allow for simultaneous signal acquisition from all channels, eliminating the deviations that arose from the sequential signal collection used in the earlier setup [21, 28]. Briefly, a femtosecond-pulsed laser with a center wavelength of 1070 nm was used to scan the sample. The generated signals were separated using dichroic mirrors and filters and detected in the epi-direction. In this study, we collected THG signals at 350-360 nm, 2PEF signals at 562-665 nm, and 3PEF signals at 380-420 nm (in an initial subset of experiments (n = 13), a filter with broadband range of 398-501 nm was used).

3D data were collected through depth scanning at 1 *μ*m intervals, covering a total depth of 20 *μ*m, with a field of view of 400 × 400 *μ*m^2^. Five regions of interest were scanned per sample, with each depth scan completed within 30 seconds.

### Data processing

Twenty depth-scanned images were saved as a single TIFF file (C × D × W × H, 3 × 20 × 1000 × 1000) using ImageJ (Fiji version 1.53). To remove the glass interface artifacts in THG channel, we applied rolling ball background subtraction with a radius of 50 pixels. Next, contrast enhancement was performed using contrast-limited adaptive histogram equalization. The data were then standardized from *v*_*i*_ to 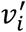, using the mean *v*_*m,i*_ and standard deviation *v*_*sd,i*_ in each channel (*i =* 1,2,3, represent the channel number), 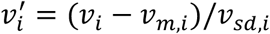, to ensure consistency across all images for further analyses.

### Model Development for Automated Cell Counting

First, we adopted the ResNet 3D-50 architecture as the foundation for our model [29]. The model consists of multiple 3D convolutional layers, pooling layers, and fully connected layers. To adapt ResNet 3D-50 for the regression task of cell counting, we replaced the original fully connected layer with a global average pooling, followed by two fully connected layers: the first with 32 units and the second with 4 units, and subsequently activated the SoftMax function to predict the cell count in four classes (Fig.2A). Each fully connected layer was regularized with a dropout rate of 0.3 to prevent overfitting. We optimized network parameters using mean absolute error (MAE) as the loss function and employed the standard stochastic gradient descent [30] method with a learning rate of 0.0012. The model was trained separately on BALF and blood fraction dataset. For the BALF dataset, the model was trained for 500 epochs, requiring approximately 20 hours. For the blood dataset, it was trained for 50 epochs. The version with the best performance on the validation set was saved for evaluation on the hold-out test set.

**Figure 2.**
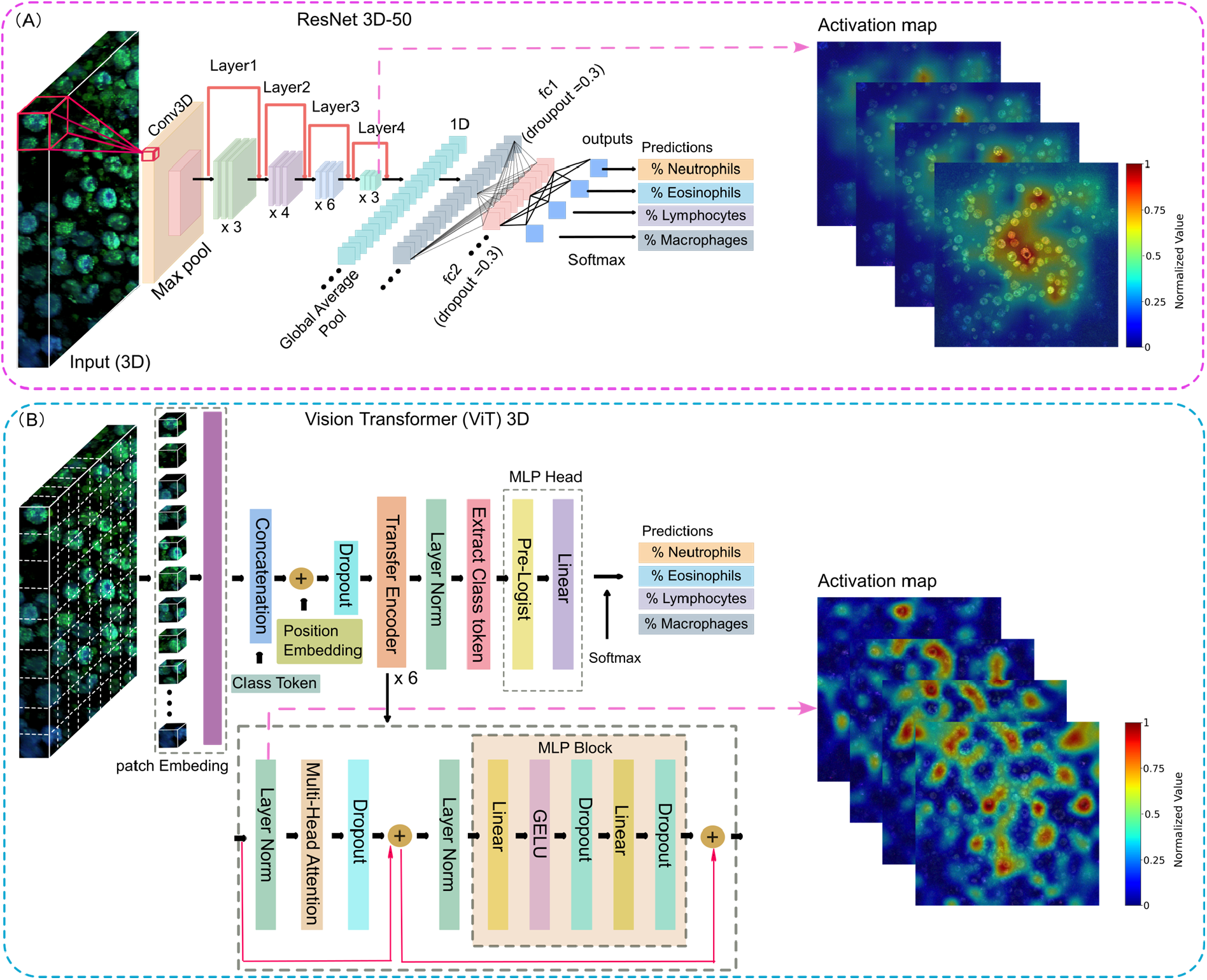
Architectural comparison of ResNet 3D-50 and ViT 3D models. The figure illustrates the key components and structural differences between the (A) ResNet 3D-50 and (B) ViT 3D architectures. ResNet 3D-50 employs 3D convolutions and residual connections to extract spatio-temporal features, while ViT 3D uses a patch-based self-attention mechanism to capture global context across the input data. Both models were utilized for 3D cell counting in this study.

Then, we applied the ViT 3D architecture to our regression task for automatic cell counting [31, 32], comparing its performance with the ResNet 3D-50 model (Fig. 2B). For this approach, the 3D images were resized to 256 × 256 × 32 × 3. The images were then divided into patches of size 16 × 16 × 2 × 3, resulting in a patch dimension of 1536 and patch number of 4096. These patches were flattened into a 1D sequence. Class tokens and positional embeddings were added to the patch sequence, which was then inputted into a standard transformer encoder. Finally, the learned class token was passed through a multi-layer perceptron (MLP) head, and a SoftMax activation was applied to produce the final output of four cell classes. The model was trained in a supervised manner to learn the distribution of cell counts, using an initial learning rate of 0.00003. A learning rate scheduler was employed, with the step size set to 1 epoch and a decay factor (gamma) of 0.7, ensuring gradual reduction of the learning rate over time. The Adam optimization algorithm was utilized to optimize the model’s performance. Like ResNet 3D-50, the ViT 3D model was trained separately on BALF and blood fraction dataset for 500 epochs (18 hours) and 50 epochs (1 hour), respectively. The version with the best performance on the validation set was saved for evaluation on the hold-out test set.

BALF and blood fraction samples were divided into training, validation, and test sets (see Table S1). Each image was labeled with the cell percentages derived from its corresponding cytological analysis. We used label smoothing strategies to solve the data distribution imbalance [33]. To enhance model robustness and generalization, we utilized two data augmentation techniques during training, including flipping and color jitter: flipping was applied to the data along x, y, and z axes, each with a 50% probability; with color jitter randomly the brightness, contrast, saturation, and hue of the image was changed by a factor of ± 20%. The hyperparameters of the ResNet 3D-50 and ViT 3D models were initially optimized using Optuna, minimizing loss value over 20 trials for 20 epochs with a median pruner strategy. The best-performing hyperparameter values were then applied for training, with additional manual fine-tuning during experiments. The final selected values are listed above. The training and validation loss curves exhibited steady convergence, indicating that the models effectively learn the patterns in the dataset (See Figure S1).

The ResNet 3D-50 and ViT 3D model were implemented in PyTorch, and the training process was conducted on a NVIDIA GeForce GTX 1080 GPU with 25 GB of RAM on the BAZIS computational cluster of the Vrije Universiteit Amsterdam.

To interpret the model’s performance, we applied Gradient-Weighted Class Activation Mapping (Grad-CAM) [34]. For the ResNet 3D-50 model, Grad-CAM computes gradients at the last convolutional layer to identify which features contribute most to the prediction of the target class (Fig. 2A). In the case of the ViT model, Grad-CAM extracts gradients at the layer normalization step of the final transformer encoder block to determine the regions driving the prediction (Fig.2B).

### Statistical Analysis

Statistical analyses were performed using Origin (Version OriginPro 2023, Academic). The results were expressed as mean ± standard deviation (SD). Bland-Altman analyses [8] were created to illustrate the average differences between the results of the cytologic analyses and models predicted results.

## Results

### Cytological analysis of leukocyte differentiation

BALF samples from ILD patients were characterized by a relative higher abundance of macrophages (34% - 88%), eosinophils (0% - 14%) and lymphocytes (1% - 50%) compared to samples from ARDS patients, with ranges representing the minimum and maximum observed values (Fig. 3A). Neutrophils were highly abundant in all BALF samples from ARDS patients (79.63% - 98.8%) (Fig. 3A). A comparison of leukocyte subpopulations in BALF and blood fractions from ARDS patients demonstrated a consistently high neutrophil percentage across both sample types, reflecting a systemic neutrophilic response in ARDS patients (Fig. 3B). Eosinophil numbers were low in BALF (0% - 0.8%) and slightly higher in blood (1.2% - 6.7%). Lymphocyte proportions were slightly higher in blood fractions (2.6% - 11.6%) compared to BALF (0.2% - 5.0%), though overall levels remained low.

**Figure 3.**
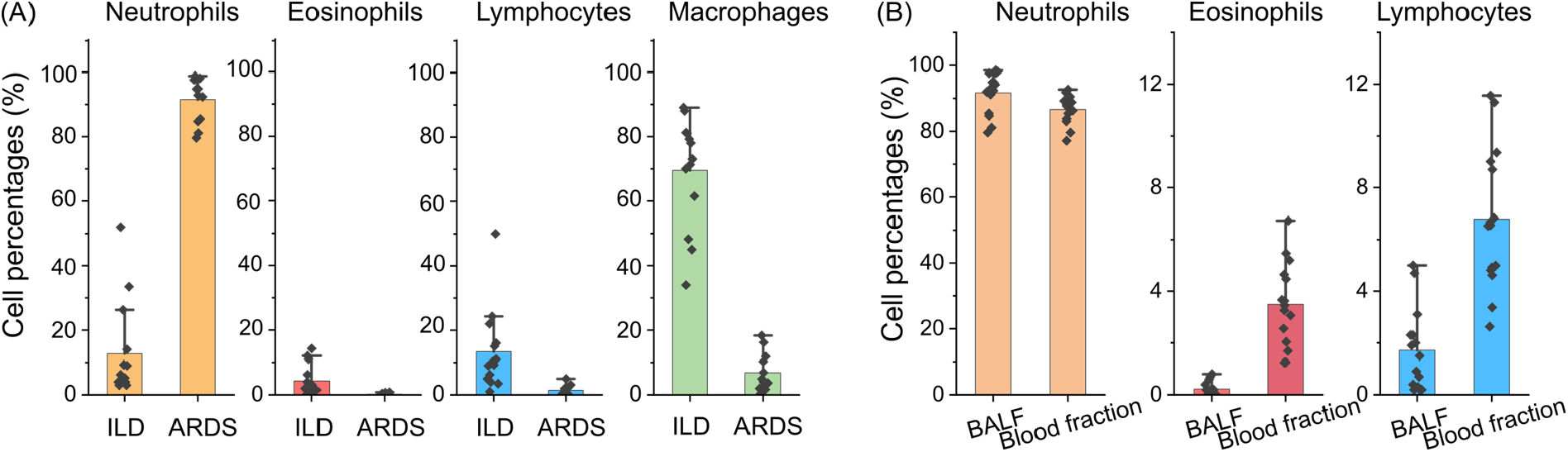
Cytological analysis of leukocyte differentiation in BALF and blood fractions of ILD and ARDS patients. (A) Comparison of leukocyte composition in BALF samples between ILD and ARDS patients. (B) Comparison of leukocyte percentages between BALF and blood fractions in ARDS patients. Shown is percentage (%) of total cell population. Data is presented as mean ± SD.

### Characterization of leukocytes via HHGM

Figure 4A presents 3D THG and MPEF volume images of BALF and blood fraction samples across three imaging channels. Randomly selected regions (boxed areas) are enlarged in Fig. 4B. Macrophages are distinguished by their larger diameter, approximately 24.8 ± 2.9µm, while neutrophils, eosinophils, and lymphocytes were smaller, with diameters ranging between 6-10µm (Fig. 4E). Neutrophils typically have a nucleus that is segmented in 3-5 connected lobes, eosinophils have a bilobed nucleus, and lymphocytes and macrophages feature a round-shaped mononuclear structure (Fig. 4B). Macrophages occasionally can appear as multinuclear cells, especially in ILD BALF samples. Additionally, neutrophils and eosinophils exhibit intracellular granules. The leukocyte characterization closely aligns with findings from a previous study [28].

**Figure 4.**
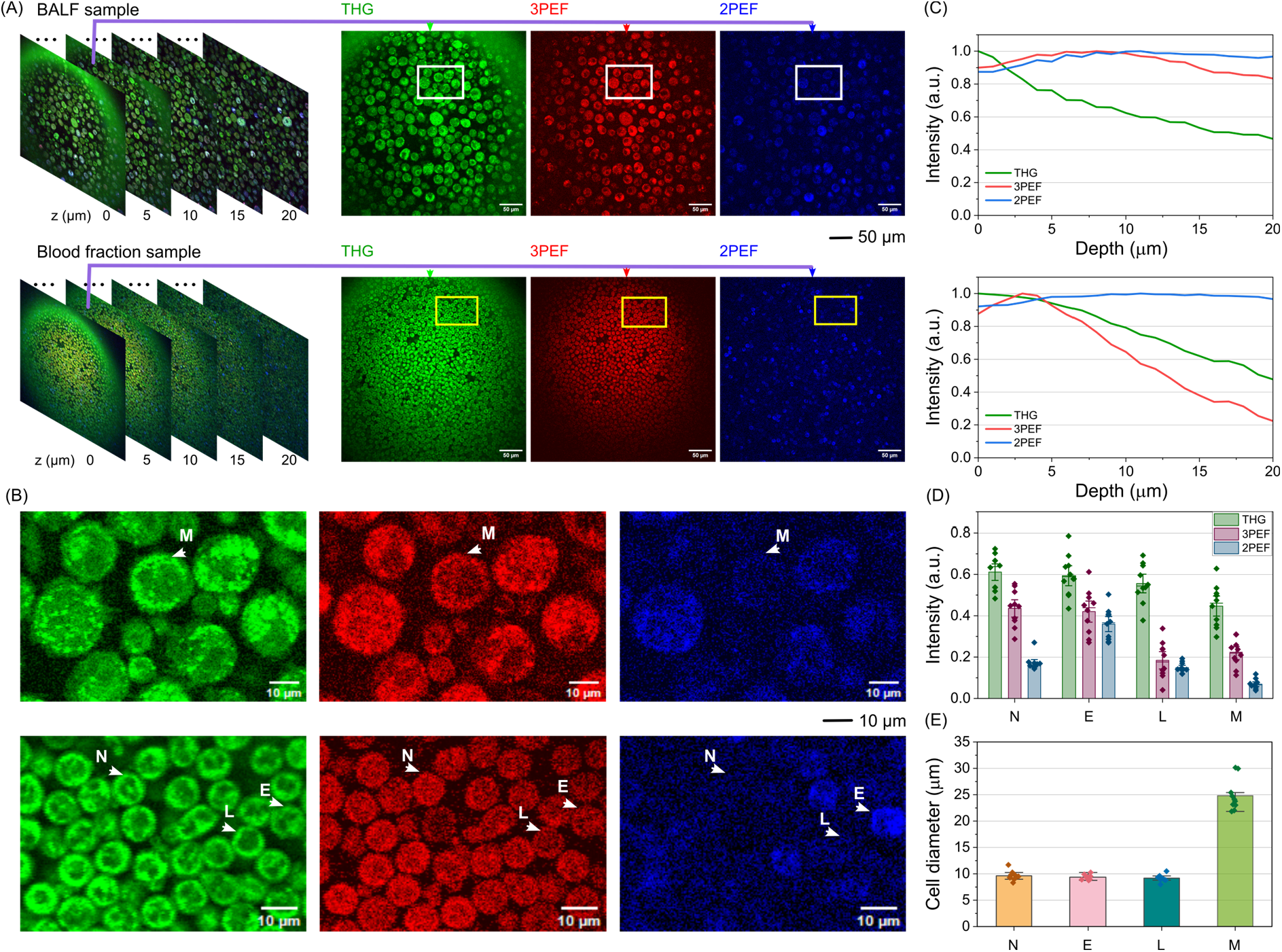
Characterization of leukocytes via HHGM. (A) 3D HHGM volume images displaying cellular structures in BALF and blood fraction samples across 3 channels. The THG signals (green) provide detailed morphological information of the cell membrane and nucleus, the 2PEF (blue) and 3PEF signals (red) originate from autofluorescence of the cytoplasm. (B) Zoomed-in views of the boxed areas in Fig. 2A, with white arrows indicating different cell types. M: macrophages, N: neutrophils, E: eosinophils, L: lymphocytes. (C) Intensity profiles of the three channels along the depth in BALF and blood fraction samples. (D) Normalized THG and 2/3PEF signal intensities, along with (E) cell diameters, for 10 randomly selected cells from each cell type.

In this study, spatial information was incorporated to enhance cell differentiation. Figure 4C displays the intensity profiles of THG, 2PEF, and 3PEF signals along the depth. THG intensity decreased with depth in both BALF and blood fraction samples. The 2PEF intensity remained stable in both sample types, while the 3PEF intensity remained stable in BALF but decreased with depth in blood samples, likely due to higher cell density in blood, which increases scattering and absorption. Eosinophils exhibited the highest 2PEF signals, while neutrophils had the highest 3PEF signals among all cell types (Fig. 4D).

### Comparison of cytology results and model predictions

The ResNet 3D-50 model predicted percentages of all cell types in BALF samples with an average accuracy >97%, closely matching the cytology results (Fig. 5A). The ViT model shows comparable performance, with average accuracy >96% (Fig. 5A). Compared to the previous study using 2D HHGM data [28], the model’s accuracy improved from 90% to 96%.

**Figure 5.**
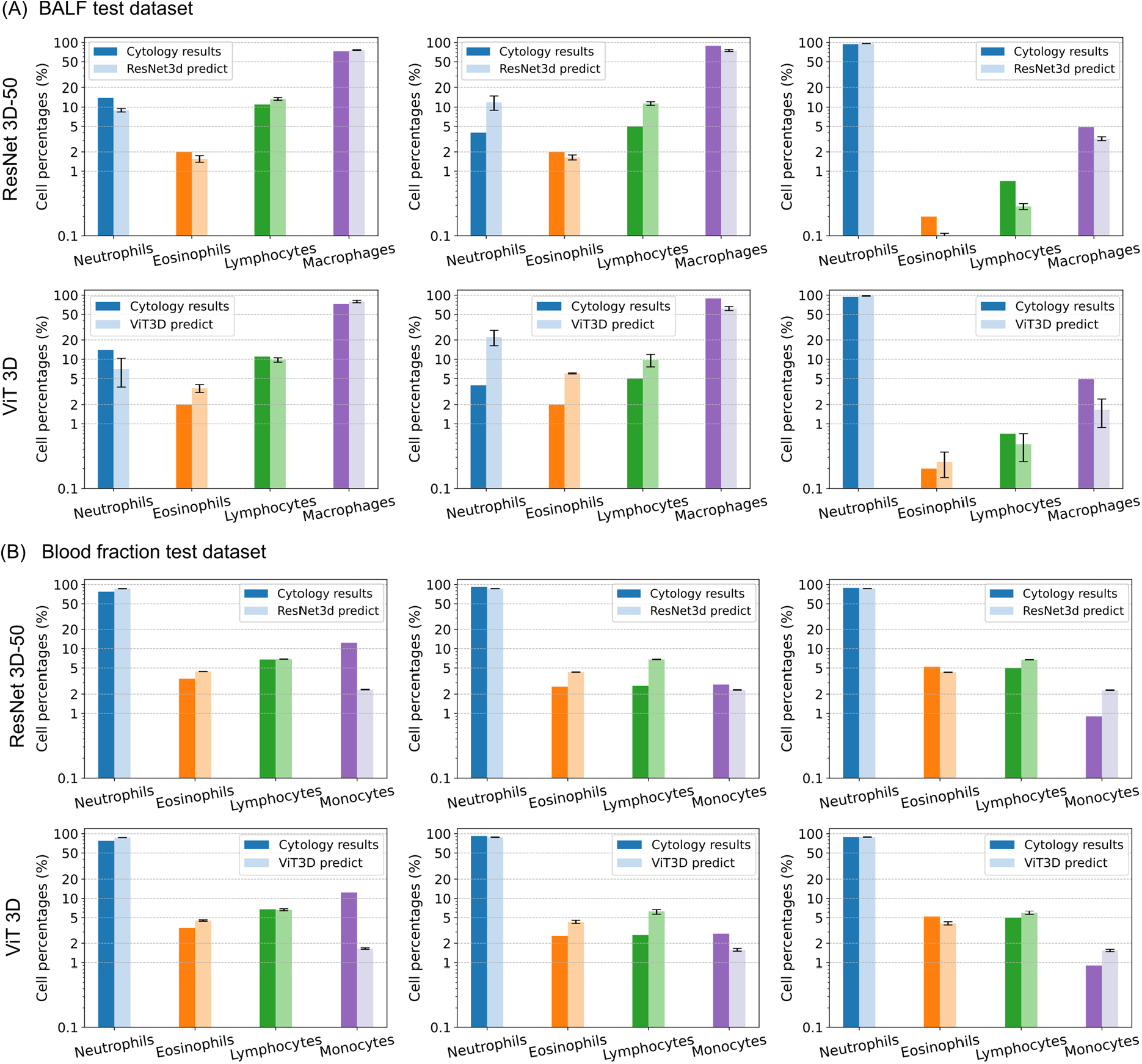
Comparison of cytology results and model predictions. (A) Predicted leukocyte compositions by ResNet 3D-50 and ViT 3D models vs. ground truth cytology results for BALF samples in test dataset. (B) Same comparison for blood fraction samples in test dataset. Dark colors represent cytology results, light colors indicate model predictions.

For the blood fraction dataset, ResNet 3D-50 predicted neutrophil percentages with high accuracy but showed slight underestimations for monocytes (Fig. 5B). Eosinophil and lymphocyte predictions were consistent with the cytology results, though minor discrepancies were observed. The average accuracy of the ResNet 3D-50 model on the blood fraction test dataset is >97%. The ViT 3D predictions consistently captured the overall trends, with prediction values aligning closely with the cytology results and an average accuracy >96%.

Bland-Altman analyses showed that the mean differences between cytology and ResNet 3D model predictions were <2% for all BALF and blood fraction leukocytes subpopulations, while those for the ViT 3D model were <3% (Fig. 6). Detailed difference of subpopulations is shown in Table 2. Moreover, the 95% confidence interval (CI) for the differences between cytology and ResNet 3D model was within the 13% margin for all leukocyte subpopulations, compared to a 25% margin for the ViT 3D model. The summary of 95% CI of all leukocyte subpopulations are summarized in Table 3.

**Table 2.** Summary of the difference of subpopulations between cytologic analyses and model predictions.

| Mean Differences |  | Neutrophils | Eosinophils | Lymphocytes | Macrophages | Monocytes |
| --- | --- | --- | --- | --- | --- | --- |
| BAL | Cytology – ResNet 3D-50 | -1.7% | 0.2% | -0.1% | 1.6% | - |
|  | Cytology – ViT3D | -2.0% | -0.9% | 0.3% | 2.6% | - |
| Blood | Cytology – ResNet 3D-50 | -1.1% | -0.6% | -0.3% | - | 1.9% |
|  | Cytology – ViT3D | -2.8% | -0.4% | 0.5% | - | 2.7% |

**Table 3.** Summary of the lower and upper agreement limit of subpopulations between cytologic analyses and model predictions.

| Lower and upper agreement limit |  | Neutrophils | Eosinophils | Lymphocytes | Macrophages | Monocytes |
| --- | --- | --- | --- | --- | --- | --- |
| BAL | Cytology & ResNet 3D-50 | -11.3%,<br>7.9% | -0.3%,<br>0.7% | -6.6%, 6.4% | -9.7%, 12.9% | - |
|  | Cytology & ViT3D | -18.7%,<br>14.7% | -3.7%,<br>1.9% | -4.5%, 5.0% | -19.1%,<br>24.4% | - |
| Blood | Cytology & ResNet 3D-50 | -12.0%,<br>9.8% | -2.5%,<br>1.3% | -6.9%, 6.4% | - | -7.6%,<br>11.5% |
|  | Cytology & ViT3D | -13.7%,<br>8.0% | -2.6%,<br>1.8% | -6.9%, 8.0% | - | -6.7%.<br>12.1% |

**Figure 6.**
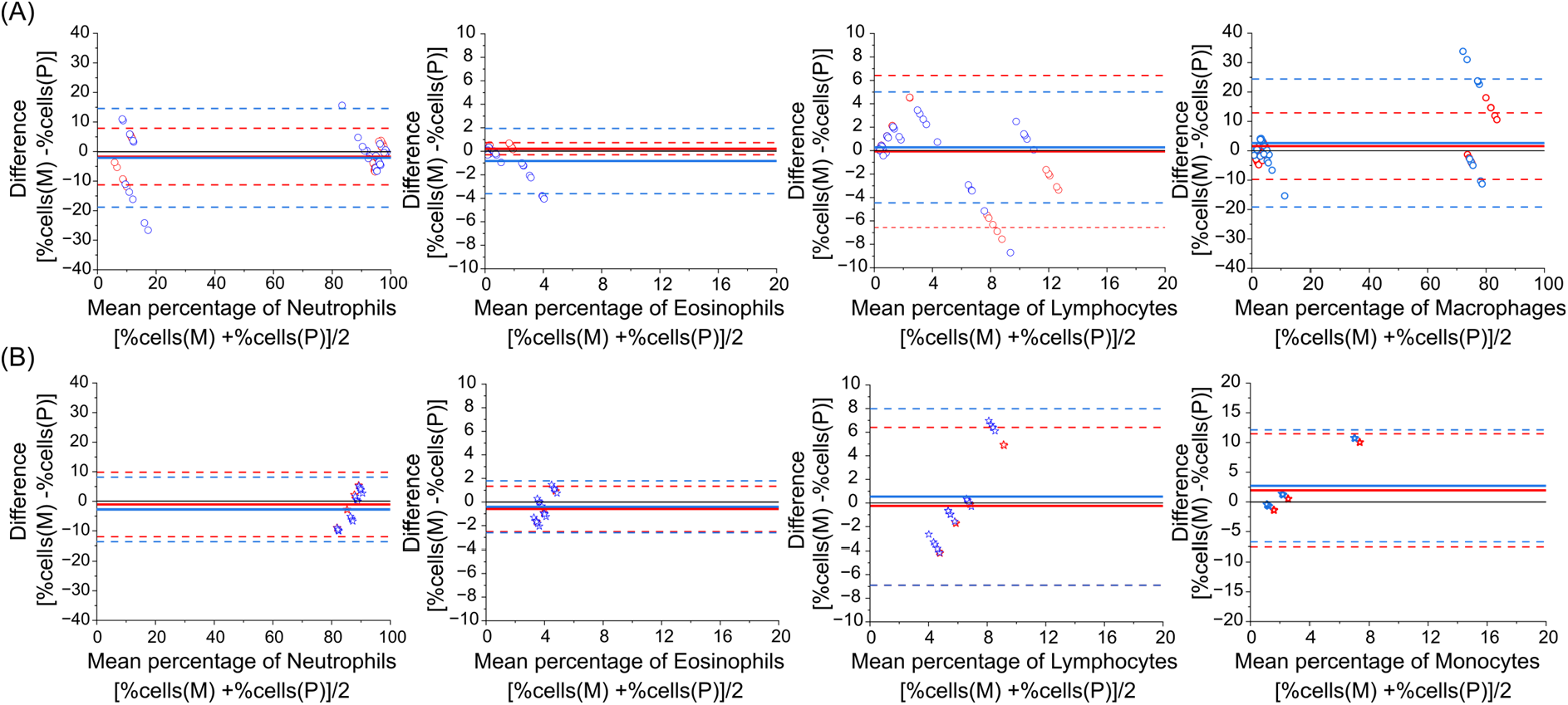
Bland-Altman analysis of cytologic and model predicted leukocyte percentages. The plots display the difference in cell percentage (cytology - prediction) on the y-axis against the mean cell percentage from both methods on the x-axis. Red represent ResNet 3D-50 predictions, and blue denote ViT 3D results. Horizontal thick lines represent average difference, dashed lines represent 95% CI of lower and upper agreement limit. (A) shows data for BAL samples (circles), and (B) shows data for blood fraction samples (stars).

## Discussion

Our findings demonstrate that our deep learning framework achieves over 96% accuracy in quantifying leukocyte composition in BALF and blood fraction samples. Validation in independent ILD and ARDS cohorts highlights its potential to streamline cell counting in diverse diagnostic settings. Whether the approach is also suitable to analyze other types of samples like bronchial washes, sputum samples, or cerebrospinal fluid for example will need further investigation.

Incorporating 3D spatial information from label-free HHGM significantly improved differentiation accuracy compared to 2D methods, achieving well concordance with gold-standard cytologic analyses (mean difference <3%; see Table 2). The ResNet 3D–50 model slightly outperformed the ViT 3D model. Importantly, high accuracy was maintained despite training on limited, image-level annotations, avoiding labor-intensive 3D cell border annotation, highlighting their efficiency and scalability. While class imbalance in the current dataset may limit generalizability for rare cell populations, future multi-center studies with larger datasets will address this limitation and further validate clinical applicability.

We approached multi-class cell counting as a regression task rather than a classification task. Although multi-class classification frameworks are commonly used [35-37], treating cell counting as a regression task allows the model to predict continuous values for cell percentages. This approach aligns better with the nature of cell counting, where the distance between different cell counts is critical—for instance, counts of 25 and 26 are closer than counts of 25 and 53 [27]. However, the regression framework’s focus on global counts (without spatial localization) introduces a trade-off, potentially causing false positives. Future integration of density regression models [26], which jointly predict counts and centroid coordinates, could enhance performance while retaining computational efficiency.

Differentiation of cells present in body fluids like blood and BALF are frequently used in both research and clinical setting. An important example is the use of BALF analysis in the diagnosis of ILDs. Our approach using HHGM images and deep learning analysis offers significant benefits in terms of cost-effectiveness, low expertise requirement, field portability, and rapid turnaround time compared to the alternatives (cytospin [38] or flow cytometry [8, 39]). Indeed, it generates 3D volumetric images with detailed cell morphology and estimates leukocyte composition percentages within minutes. A limitation compared to flow cytometry-based analyses may be that our approach does not provide information about cell marker expression due to limited resolution. Unlike microscopy or flow cytometry, our method eliminates the need for staining, specialized facilities, or extensive training, offering field portability and near-instant results. Furthermore, the system requires only a small sample volume, making it particularly suitable for field applications. Thus, these advantages position the proposed system as a promising diagnostic tool, especially in resource-limited settings where access to conventional medical diagnostics may be limited.

## Conclusions

The combination of the HHGM and deep learning presents a promising approach for differentiating leukocytes in fresh, label-free BALF and blood fraction samples. This technique has the potential to accelerate the diagnostic process while reducing costs, needed expertise and overall workload.

## Supporting information

Supplementary Information added and link to the file on the preprint site

## Abbreviations

ARDS: acute respiratory distress syndrome
BALF: broncho-alveolar lavage fluid
CI: confidence interval
DQ: Diff-Quick
HHGM: higher harmonic generation microscopy
ILD: interstitial lung disease
MAE: mean absolute error
MLP: multi-layer perceptron
MPEF: multiphoton excited autofluorescence
SD: standard deviation
THG: third harmonic generation
2PEF: two-photon excited autofluorescence
3PEF: three-photon excited autofluorescence
ViT: Vision transformer.

## Supplementary Information

The supplementary information can be found in Supplementary 1.

## Declarations

## Ethics approval and consent to participate

For ARDS patients, informed consent was deferred until ICU discharge. Patients were prospectively included if they provided written informed consent, or if neither they nor their relatives opted out, or if the patient had died. For ILD patients, all patients provided written informed consent.

The studies were approved by the Medical Research Board under protocol numbers 2020.213 (BALI biobank) and 2022.0507 (Amsterdam Interstitial Lung Disease biobank). The research was conducted in accordance with the Netherlands Code of Conduct for Research Integrity and the ethical guidelines of the Declaration of Helsinki.

## Consent for publication

Not applicable

## Availability of data and materials

The datasets used during this study are available from the corresponding author on reasonable request.

## Competing interests

M.L.G has an indirect interest in Flash Pathology B.V., the other authors declare no competing interests.

## Funding

M.Z was funded from a scholarship from China Scholarship Council (CSC202006970016).

## Author’s contributions

M.L.G and J.W.D designed research; M.Z performed the HHGM experimental work; T.D, S.Z, L.S.B, H.B.v.d.H, A.D, I.A.S processed the clinical samples; T.D performed the cytologic analyses; M.Z performed the computational work, P.J.G supervised the computational work; all authors contributed to data interpretation; M.Z wrote the manuscript, all authors read and edited the manuscript.

## Acknowledgements

We are grateful to Frank van Mourik from Flash Pathology B.V. for his technical support with the multiphoton microscopy setup. We are grateful to the collaborators of the Amsterdam UMC BALI study group and this paper is published on their behalf: Peter I. Bonta, Lieuwe D.J. Bos, Justin de Brabander, Leo Heunks, Erik Michels, Esther J. Nossent, Tom van der Poll and Alexander P.J. Vlaar.

## Notes

### Competing Interest Statement

M.G. declares to have financial and non-financial interest in Flash Pathology B.V. However, Flash Pathology B.V. was not involved in the design of the study or analysis of the data. The other authors declare no competing interests.

