## Supplementary Information added and link to the file on the preprint site for "Deep Learning-Based 3D Leukocyte Differentiation Using Label-Free Higher Harmonic Generation Microscopy"

**Table S1. Data overview for training, validation, and test sets [a]**

| Sample type | Patient condition | Training | | Validation | | Test | |
|---|---|---|---|---|---|---|---|
| | | sample | 3D stacks | sample | 3D stacks | sample | 3D stacks |
| BALF | ILD | 9 | 45 | 3 | 15 | 2 | 10 |
| | ARDS | 9 | 45 | 2 | 10 | 5 | 25 |
| Blood fraction [b] | ARDS | 8 | 40 | 3 | 15 | 4 | 20 |
| Total | | 26 | 130 | 8 | 40 | 11 | 55 |

[a]: We initiated model training after collecting 34 samples, with 80% randomly assigned to the training set and 20% to the validation set. Subsequently, we acquired an additional subset for the test set, comprising approximately 24% of the entire dataset. Each sample, consisting of five 3D stack images, was assigned entirely to one subsets to prevent data leakage. Notably, three ARDS patients underwent multiple bronchoscopies and blood collections at different time points (two patients sampled twice, one sampled three times), each collection was treated as a distinct sample due to unique cytospin analysis results.

[b]: one blood fraction sample was excluded due to different cell counter processing.

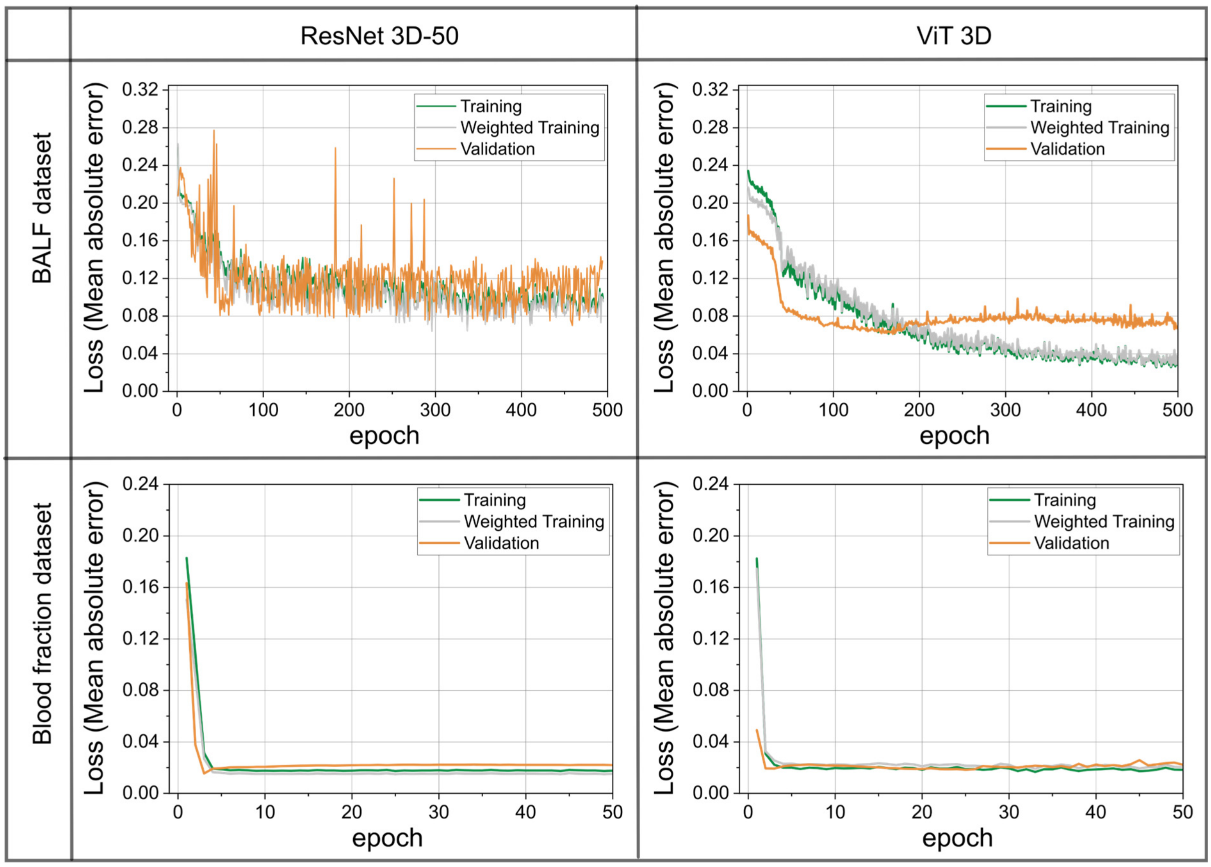


**Fig.S1. Training and validation loss curves for the optimal ResNet 3D-50 and ViT 3D models on the BALF and blood fraction datasets, selected from multiple experimental trials for lowest final validation loss**. The weighted training loss, computed under label smoothing, addressed data imbalance. ResNet 3D-50 showed steady convergence, with validation loss stabilizing at ~0.08–0.1 MAE, while ViT 3D exhibited more stable convergence, reaching ~0.06–0.08 MAE within 100 epochs. For the blood fraction dataset, both models converged within 50 epochs, with training and validation losses stabilizing at ~0.02–0.03 MAE.
